# Variation in thermal physiology can drive the temperature-dependence of microbial community richness

**DOI:** 10.1101/2022.10.28.514215

**Authors:** Tom Clegg, Samraat Pawar

**Affiliations:** The Georgina Mace Centre for the Living Planet, Department of Life Sciences, Silwood Park Campus, Imperial College London, Buckhurst Road, Ascot, SL5 7PY, UK

**Author notes:** **For correspondence:** (TC).

## Abstract

Predicting how species diversity changes along environmental gradients is an enduring problem in ecology. Current theories cannot explain the observation that microbial taxonomic richness can show positive, unimodal, as well as negative diversity-temperature gradients. Here we derive a general empirically-grounded theory that can explain this phenomenon by linking microbial species richness in local communities to variation in their temperature-driven competitive interaction and growth rates. It predicts that richness depends on variation in shape of the thermal performance curves of these metabolic traits across species in the community. Specifically, the shape of the microbial community temperature-richness relationship depends on how the strength of competition across the community and the degree of variation in growth rates changes across temperature. These in turn can be predicted from the variation in thermal performance across the community. We show that empirical variation in the thermal performance curves of metabolic traits across extant bacterial taxa is indeed sufficient to generate the variety of community-level temperature-richness responses observed in the real world. Our results provide a new mechanism that can help explain temperature-diversity gradients in microbial communities, and provide a quantitative framework for interlinking variation in the thermal physiology of microbial species to their community-level diversity.

## Introduction

The effect of temperature on biodiversity has long been a topic of interest in ecology. Starting with the pioneering work of Alexander von Humboldt who in the 19th century identified temperature as a major environmental driver of plant richness along elevational gradients in the Andes (***von Humboldt and Bonpland, 2013***), temperature has been recognized as a key driver of the geographical gradients in taxonomic richness seen across practically all organismal groups (***Rohde, 1992***; ***Gaston, 2000***). In recent years, the relationship between species richness in microbial communities and temperature in particular has become a topic of particular interest, in concert with an increase in awareness of the importance of these communities to ecosystem functioning (***Schimel, 2013***; ***Graham et al., 2016***; ***Antwis et al., 2017***), and new DNA sequencing technologies that allow community “snapshots” to be characterised with relative ease (***Zimmerman et al., 2014***). Studies on microbial community richness, most often measured in numbers of OTUs (operational taxonomic units), have generally found varying responses to changes in environmental temperature. For example, while ***Zhou et al. (2016***) found that soil microbe richness increased across a continental temperature gradient in North America, others have found unimodal responses (richness peaking at intermediate temperatures) in soils as well as other environments (***Milici et al., 2016***; ***Sharp et al., 2014***; ***Thompson et al., 2017***). Indeed, as demonstrated by the data-synthesis by ***Hendershot et al. (2017***), the temperature responses of microbial richness or diversity are “consistently inconsistent”, with no single pattern in terms of shape (monotonic or unimodal) or direction (positive or negative) dominating.

Currently, two mechanistic explanations exist relevant to microbial temperature-richness gradients over space, both focusing on energy availability in the environment. First is the metabolic theory of biodiversity (MTB) (***Allen et al., 2002***), which predicts a monotonic increase in species richness with temperature because of increasing cellular kinetic energy at higher temperatures. This allows more individuals to survive in a given community which in turn supports higher species richness. However, the MTB relies on two strong assumptions: energy equivalence (all populations have the same rate of energy use), and a single temperature dependence across taxa (the “Universal Temperature Dependence” or “UTD”). While the validity of energy equivalence in the microbial world remains untested to the best of our knowledge, the emergence of extensive evidence demonstrating significant functional variation in thermal sensitivities across the tree of life indicates that the UTD assumption is at best an approximation (***Smith et al., 2019***; ***Dell et al., 2011***; ***Kontopoulos et al., 2020***). Furthermore, the MTB does not consider resource-mediated species interactions (especially, competition) and can only predict a positive-monotonic temperature-diversity response, which cannot account for the commonly observed unimodality. A variant of the MTB model was later introduced by ***Stegen et al. (2012***) that can generate unimodal temperature-diversity gradients by accounting for muti-trophic species interactions and resource limitation. This model relaxes the energy equivalence assumption, retains the UTD assumption, and adds two new ones: body size variation as a source of diversity, and a temperature-dependent resource supply. Whilst the temperature dependence of resource supply will undoubtedly be an important factor in microbial communities the UTD assumption, along with body size scaling are of questionable relevance to microbial communities, where body (cell) size is not expected to be as big a source of variation in functional diversity in microbes as it is in food webs (***Westoby et al., 2021***).

Second, the metabolic niche hypothesis (***Sharp et al., 2014***), offers an alternative explanation for microbial temperature-diversity gradients. It posits that there are more energetically-viable ways to make a living at intermediate (non-extreme) temperatures, thereby allowing greater species co-existence and ultimately yielding a unimodal temperature-diversity response (***Clarke and Gaston, 2006***; ***Marsland et al., 2020***). However, the mathematical model used to explore this hypothesis (***Marsland et al., 2020***) does not explicitly link the temperature dependence of ecologically relevant metabolic traits to coexistence and species diversity, and furthermore, much like the MTB, cannot explain the full spectrum of shapes of the the temperature-diversity responses of microbes, where unimodality is by no means a universal rule.

A key aspect missing from these current explanations is the significant variation that exists in thermal responses of key metabolic traits such as growth and respiration rate across microbial taxa (***Smith et al., 2019, 2021***; ***Kontopoulos et al., 2020***). This variation is expected to be important in determining the responses of microbial communities to temperature through two mechanisms. First, the nonlinear thermal responses of individual- and population-level metabolic rates mean that inter-specific variation in thermal sensitivity may result in significant changes in realised trait-value distributions at different temperatures. In such cases considering only the average of thermal responses is likely to be a poor approximation (***Savage, 2004***). Second, differences in thermal responses of traits of interacting populations (“physiological mismatches”) can have non-trivial effects on community level dynamics, including coexistence (***Dell et al., 2014***; ***Bestion et al., 2018***; ***García et al., In press***).

In this paper, we derive a new theory that predicts the response of microbial community richness to temperature while accounting for variation in thermal sensitivity of metabolic traits across populations. We focus on competitive interactions, which strongly limit diversity in microbial communities in particular (***Marsland et al., 2019***; ***Goldford et al., 2018***; ***Ratzke et al., 2020***; ***Lechón et al., 2021***). Using a general model, we first derive a mathematical expression that links the distribution of population thermal performance curves to the number of species that can feasibly coexist within a community. Then, using empirical data to parameterise the model, we ask whether the extant variation in thermal responses of bacterial metabolic traits is sufficient to be a key driver of patterns of species richness across temperature gradients in the real world. We use extensive numerical simulations to show that our analytical results are both robust and general.

## Results

### Theory

We link the effects of a given community-level distribution of two key metabolic traits—maximal population growth rate *r*(*T*) and pairwise interaction strengths *a*_*ij*_ (*T*) —to the probability of feasibility (*P*_*feas*_): the probability that the community will support all species’ populations at non-zero abundance at equilibrium. Feasibility is a necessary condition for stable population coexistence (***Grilli et al., 2017***; ***Dougoud et al., 2018***) and thus determines the maximum number of species in any given community. We then determine how temperature, acting through these two traits, changes *P*_*feas*_ and thus richness. ***Figure 1*** provides an overview of the theory.

**Figure 1.**
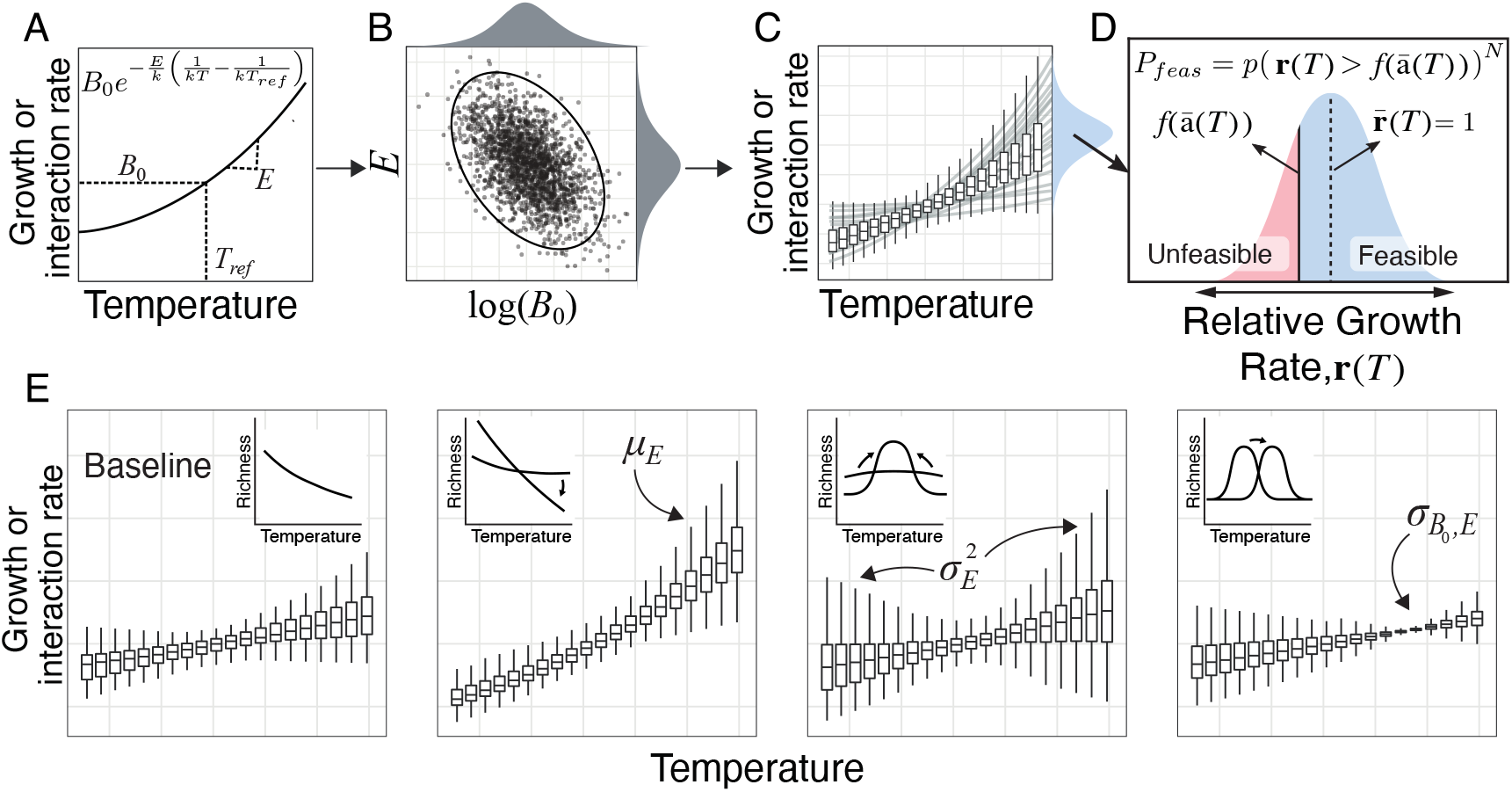
How thermal physiology constrains microbial community species richness. **(A)** Trait values increase with temperature following the Boltzmann-Arrhenius model (***Equation 14***) governed by two parameters: *B*_0_ - trait value at a reference temperature *T*_*ref*_ and *E* - thermal sensitivity. **(B)** The joint distribution of *E* and log(*B*_0_) (here with empirically-realistic negative covariance) determines how trait distributions vary across temperatures **(B)**. **(D)** The distribution of trait values in turn determines the probability of feasibility *P*_*feas*_ (and thus richness; ***Equation 1***) in terms of the emergent community-level relative growth rate **r** and mean interaction strength (*ā*). *P*_*feas*_ is determined by the proportion of relative growth rates (blue shaded area) that are greater than the bound set by mean interaction strength (solid black line). Populations with relative growth rates below this bound (red shaded area) are unfeasible (cannot persist in the community). All else being equal, the size of the unfeasible region (*P*_*f eas*_), i.e., richness, decreases with increasing variance in the growth rate distribution (Var(**r**)) and increasing interaction strengths (which shifts the *f* (*ā* (*T*)) bound upwards). **(E)** The effects of varying different aspects of the joint distribution of *B*_0_ and *E* on the emergent trait distribution across temperatures. Each panel shows the effect of altering the labeled parameter relative to the baseline case (far left), with inset plots showing the effect on the resulting temperature-richness relationship.

#### Community-level trait distributions determine species richness

We first show that the probability of community feasibility *P*_*feas*_ is given by (Methods):

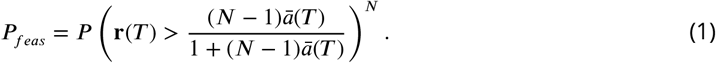

Here, *N* is species richness, 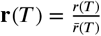 is the relative growth rate (i.e., its value relative to the average *r* (*T*) of all *N* populations), and 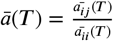 is the average of interspecific competitive interaction strength *ā*_*ij*_ (*T*) normalised by the average intraspecific interaction strength *ā*_*ii*_(*T*). The inequality inside the brackets represents the probability that a given population is feasible (i.e. has non-zero biomass) with the *N*th power term representing the fact that all populations must meet this criteria for a community to be feasible. Note that this expression for *P*_*feas*_ is in terms of the two community-level parameters **r**(*T*) and *ā* (*T*). From this point we make the important distinction between these community-level properties which emerge from the population-level traits, individual population growth rates *r*_*i*_(*T*)s and interaction coefficients *a*_*ii*_(*T*)s, which they are based on.

***Equation 1*** states that community feasibility is maximised when the negative effects of interspecific competition on each population (right hand side of the inequality) do not outweigh their relative growth rates (**r**(*T*); ***Figure 1***D). Therefore, *P*_*feas*_ declines with increasing *N*, placing an upper bound on richness, for two reasons. First, assuming that the average strength of individual competitive interactions is constant, the addition of new species to a community will result in the overall strength ((*N*−1) *ā* (*T*) term) of competition increasing, reducing population abundances. This reduces the chance that the inequality in ***Equation 1*** holds for all species, thus reducing *P*_*feas*_. Second, *P*_*feas*_ will fall as *N* increases also because it becomes increasingly unlikely that the inequality in the brackets holds across all *N* species. This is effectively the same as saying that if we were to randomly sample a set of species to create a community community the chance that some might have relatively low **r**(*T*) values increases with *N*. Ecologically, this means greater community feasibility (higher richness) can be achieved through two corresponding counter-mechanisms. First, the overall strength of competition of a community can be reduced through mechanisms such as resource partitioning. This lowers *ā* (*T*) and thus the magnitude of the RHS of ***Equation 10***. Second, variation in growth rates can be reduced so that species’ **r**(*T*) values are relatively similar. This reduces the probability that a species’ **r**(*T*) value falls below the bound set by competitive interactions (i.e., that its relative growth rate is to low to endure the negative effects of competition).

#### Variation in thermal physiology determines temperature-specific trait distributions

We next decompose the above feasibility condition into the (temperature-dependent) contributions of the population-level traits *r*(*T*) and *a*_*ij*_ (*T*) which in turn determine **r**(*T*) and *ā* (*T*) (see Methods). We use the Boltzmann-Arrhenius equation to represent the exponential-like thermal performance curves (TPCs) of these population-level traits (***Figure 1***A). The distribution of trait values at different temperatures will be determined by how the two parameters of the Boltzmann-Arrhenius TPC, *B*_0_ and *E*, co-vary across species. Specifically, we show that the temperature-dependent distribution of either trait is given by:

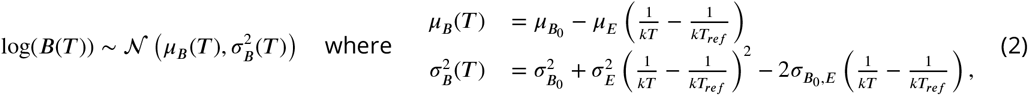

where *B*(*T*) denotes either trait’s value (*r*(*T*) or *a*_*ij*_ (*T*)), *k* is the Boltzmann constant (eV), *T* is temperature (Kelvin), and *T*_*ref*_ is the reference temperature (Kelvin). ***Equation 2*** states that the distribution of trait values *B*(*T*)’s follows a log-normal distribution with temperature-specific mean and variance, which are themselves determined by the means 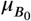 and *μ*_*E*_, variances 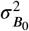 and 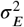 and covariance 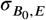 of the two TPC parameters (***Figure 1***B), as illustrated in ***Figure 1***E. Specifically: (i) A higher mean thermal sensitivity (*μ*_*E*_) across species in the community increases not just the mean trait value with temperature but also its variance; (ii) Increasing variance in thermal sensitivity 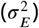 increases trait variance at extreme temperatures. In the absence of covariance this occurs either side of the reference temperature *T*_*ref*_; (iii) The covariance 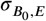 determines the temperature where the lowest trait variance occurs, with negative covariance values shifting this point towards warmer temperatures. We henceforth focus on the case of negative covariance 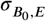 because this is the pattern observed in empirical TPCs (further Results below), consistent with the existence of a trade-off between trait performance and thermal sensitivity (a generalist-specialist tradeoff; ***Huey and Hertz*** (***1984***); ***Kontopoulos et al. (2020***)).

#### Temperature determines richness by altering community-level trait distributions

Applying ***Equation 2*** to the two traits that determine feasibility (and thus richness), 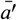 and **r**, we obtain:

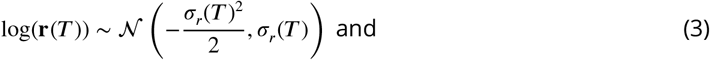

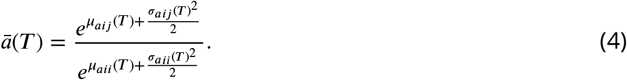

That is, the distribution of relative growth rate **r** at a given temperature is determined solely by the variance in *r*, while mean competitive interaction strength *ā* at that temperature is determined by both the mean and variance of inter- and intraspecific interaction strength *a*_*ij*_ and *a*_*ii*_. By substituting the expressions ***Equation 3*** and ***Equation 4*** into the feasibility condition ***Equation 1***, we can now predict the temperature-richness relationship in terms of the variation in thermal physiology (trait TPCs) across species in the community (encoded by 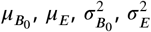 and 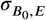; ***Figure 1***E, ***Figure 2***). Note that this step requires that we assume that the distribution of **r** is well characterised by ***Equation 2***. It is possible that realised community distributions will deviate from this if processes such as species-sorting select for species such that the distribution of **r** changes (see below).

**Figure 2.**
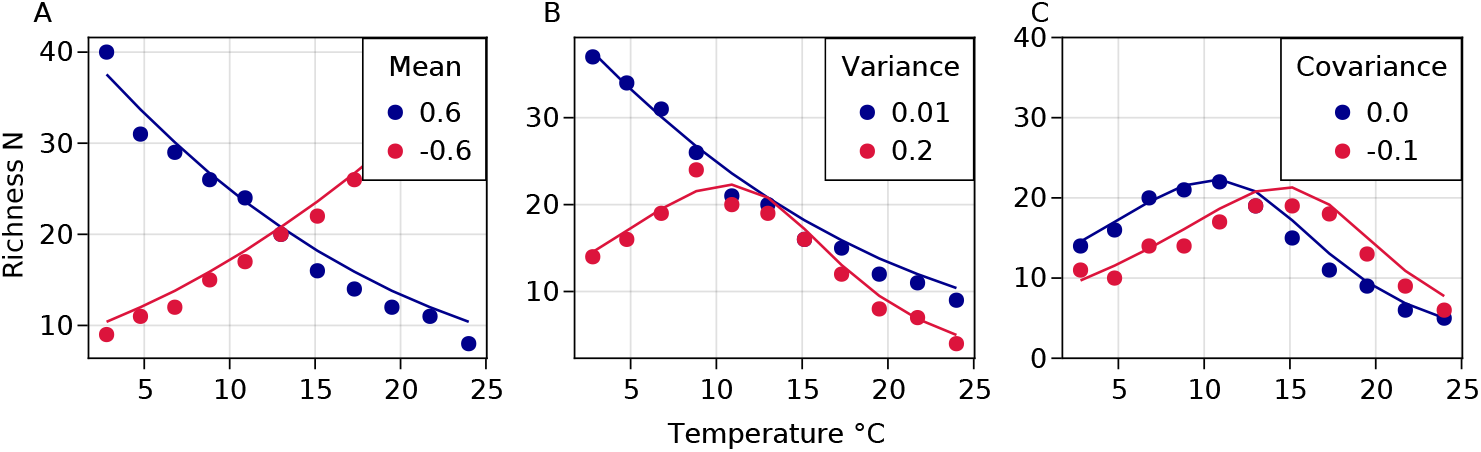
The effect of variation in trait TPCs on the temperature-richness relationship in competitive microbial communities. The analytical predictions (solid lines) are juxtaposed on the richness reached in the numerical simulations (dots). **(A)** Mean thermal sensitivity of interactions *ā* determines the direction and steepness of the temperature-richness relationship. **(B)** Increasing variance of thermal sensitivity increases unimodality. **(C)** Negative covariance between *B*_0_ and *E* shifts the peak of richness to higher temperatures. Parameter values used were: 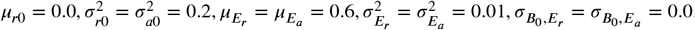.

This leads to three key insights. Firstly, the average thermal sensitivity *μ*_*E*_ will determine the rate at which richness exponentially changes with temperature (***Figure 1***E 2nd panel, ***Figure 2***A). The response of effective competition *ā* to temperature is determined primarily by the difference between the average thermal sensitivity of inter- and intraspecific interactions 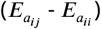 which we assume will both have a positive temperature dependence. If interspecific interactions are more sensitive 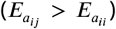 then *ā* will increase with temperature resulting in the co-existence of fewer populations and lower richness. If intraspecific interactions are more sensitive 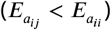 then the effective strength of competition will decrease with temperature thus leading to more populations coexisting. Note that in the case where they have the same temperature dependence the strength of effective competition will be constant over temperature and richness will be determined entirely by **r**(*T*). Secondly, increasing variance in thermal sensitivity 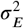 will result increased unimodality in the thermal response of richness (***Figure 2***B), with a more pronounced peak of richness at the reference temperature (set here to 13°*C*). This peak occurs as increasing 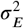 results in larger variance in **r** at extreme temperatures which means relatively fewer species are able to endure the negative effects of competition, reducing richness. Finally introducing a negative covariance between *B*0 and *E* will shift the thermal response of richness upwards towards higher temperatures, resulting in richness peaking at higher temperatures (***Figure 2***C). Overall, these predictions were matched well by the simulations using the GLV model with all parameters having the same effects predicted by the analytical theory.

### The theory holds in dynamically-assembled communities

The temperature-species richness relationship in dynamically-assembled communities also matched the theoretical predictions qualitatively (and in most cases quantitatively) ***Figure 3***. Only when varying the average thermal sensitivity did the theoretical predictions deviate from the simulations. In this case the thermal sensitivity of richness displayed a greater increase than predicted in the simulations though it still increased as qualitatively predicted ***Figure 3***B-C. This exaggerated thermal response is likely due to the effects of assembly on community trait distributions. These trait distributions change as the condition of invasiveness (i.e. the capacity for a given species invade a system) filters species with specific traits. This causes the invasion simulations to break the assumption of a constant distribution of growth rates, leading to the observed divergence in richness.

**Figure 3.**
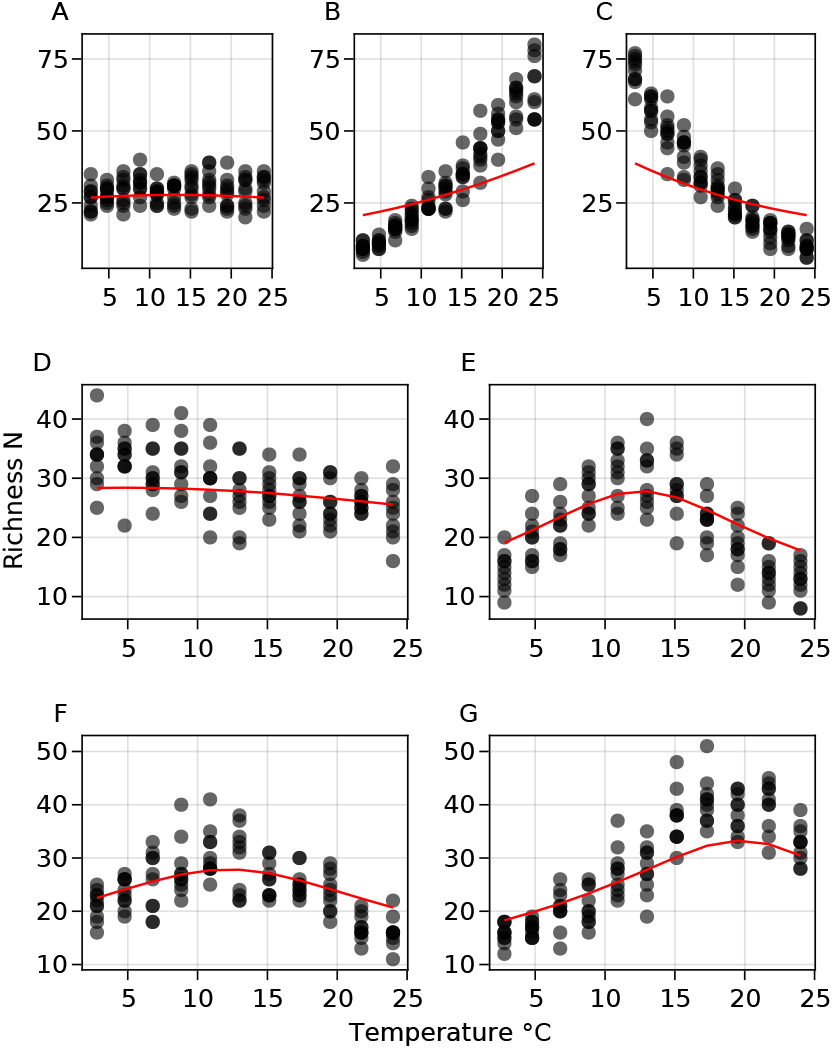
The temperature-richness relationship in dynamically-assembled microbial communities. Across all subplots, each dot represents the richness reached by a single dynamically-assembled community at the given temperature. Red lines show the predicted richness as per ***Equation 11***. (**A–C**) The thermal sensitivity of richness change with mean trait thermal sensitivity (from **A**: *μ*_*E*_ = 0, to **B**: *μ*_*E*_ = −0.6 and **C**: *μ*_*E*_ = 0.6). (**D–E**) The strength of the peak in thermal sensitivity increases as variation in sensitivity increases from **D** 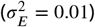 to **E** 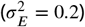 (F–G) The position of the peak shifts upwards towards higher temperatures as the covariance 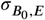 becomes more negative 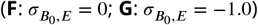. Other parameters used across all simulations are 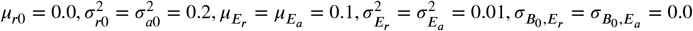.

### Real variation in thermal physiology predicts unimodal bacterial temperature-richness relationships

We next parameterised our theory with empirical data on bacterial traits to determine the temperature-richness relationship predicted under realistic levels of variation in thermal physiology ***Figure 4***. Bacterial growth rates from both experimental (***Smith et al., 2021***) and literature-synthesised (***Smith et al., 2019***) datasets showed considerable variation in TPCs thorugh the variation in both *B*_0_ and *E* and a negative covariance between log(*B*_0_) (for a *T*_*ref*_= 13° C) and *E* values (***Figure 4***A-D). Fits to the multivariate-normal distribution using MLE yielded estimates of 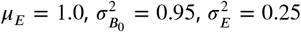 and 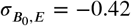 for the experimental data set and 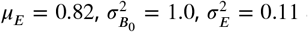 and 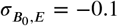 for the data-synthesis. Parameterising our theory with these values (using the same thermal response for growth rates and interactions) predicts unimodal temperature-richness responses due to this combination of variance and negative covariance ***Figure 4***E. Due to its larger variance in *E* as well as stronger negative covariance, the response based on the experimental data shows a sharper increase in richness, and peaks at a higher temperature of ∼ 20°C, than that based on the data-synthesis which has a shallower, broader temperature-richness curve peaking at ∼ 9°C.

**Figure 4.**
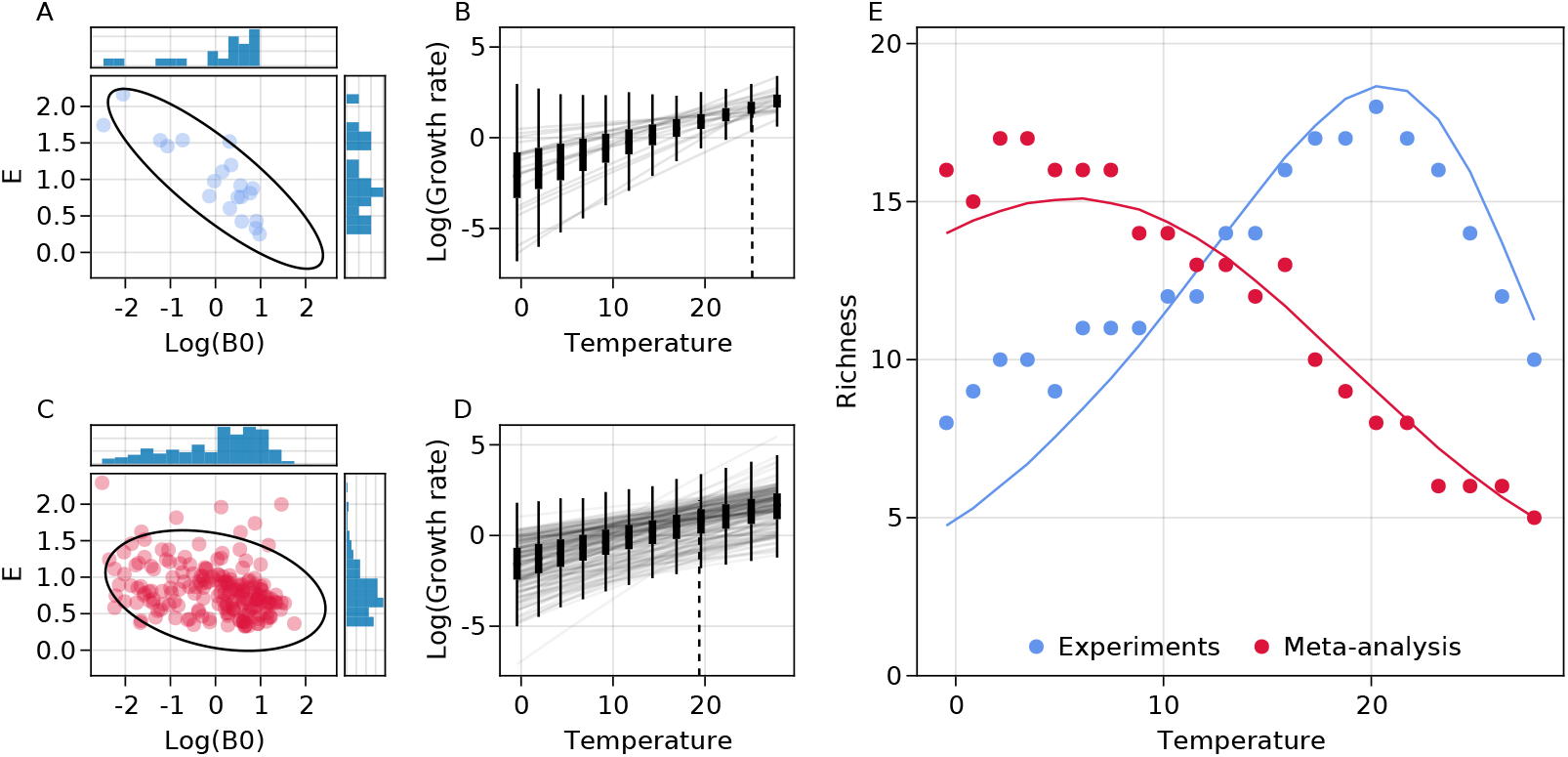
The bacterial temperature-richness relationship predicted by empirically-observed variation in thermal physiology. **(A)** The relationship between log(*B*_0_) and *E* for growth rate in the experimental TPC data from ***Smith et al. (2021***). Dots show each individual estimated *B*_0_ and *E* pairs with histograms showing their marginal distributions. Ellipses show the 95% quantiles of the fitted bivariate normal distribution. **(B)** The actual growth-rate TPCs (solid lines) from the dataset as well as the fitted trait-distributions across temperature (box-plots). The dashed line shows the point of minimum variance in growth rates which occurs towards the upper end of the temperature range. **(C-D)** Analogous plots for the dataset from the literature synthesis (***Smith et al., 2019***). **(E)** The analytically- (solid line) and simulation- (points) predicted temperature-richness curves based on the TPC variation seen in both these experimental (blue) and literature-synthesised (red) empirical data. Both are generated using the parameters from their respective fitted distributions and mean normalisation constants of 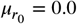 and 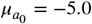. We set the normalisation constants to ensure…

## Discussion

We have investigated how variation in species-level thermal responses (TPCs) of two key metabolic traits—maximal growth and competitive interaction rate—affects the temperature dependence of species richness (the temperature-richness response) in microbial communities. We find that the shapes of the across-species distributions of thermal sensitivity (*E*) and normalisation constant (*B*_0_) parameters of the TPCs of both traits, as well as the covariance between them, are likely to be an important driver of the microbial community temperature-richness response across spatial or temporal environmental gradients in the real world.

A key new insight from our theory is that variance in thermal sensitivity of growth rate, *σ*_*E,r*′_ results in unimodality of the microbial temperature-richness curve (***Figure 2***). This is due to the quadratic temperature dependence of trait variance (***Equation 2***) and its effects on the community-level traits that determine richness (***Equation 3, Equation 4***). Furthermore, the temperature at which richness peaks in such a uninmodal response is governed by the covariance between thermal sensitivity and baseline value of growth rate (*r*_0_), with empirically-realistic negative covariance values shifting peak richness towards higher temperatures. This negative covariance is consistent with a previously identified thermal specialist-generalist tradeoff in which species tend to perform moderately across a wider temperature range (high *B*_0_ low *E*_*r*_) or better at a narrower range of temperatures at the upper end of their OTR (i.e. thermal optima; low *B*_0_ high *E*_*r*_) (***Huey and Hertz, 1984***; ***Kontopoulos et al., 2020***).

Altogether, our results reveal a new mechanism—the effect of variation in thermal physiology on competitive exclusion—by which temperature influences community richness beyond energy availability alone. We expect this mechanism to play a significant role in determining patterns of microbial richness in nature because there exists extensive variance in the thermal sensitivity *E* of metabolic traits across the microbial tree of life, as well as negative covariance between this parameter and the normalisation constant (*B*_0_) (***Kontopoulos et al., 2020***; ***Smith et al., 2019***). Our theory predicts that the decline of species richness above intermediate temperatures occurs because variation in growth rate across populations reduces the number of species that are able to persist under competitive pressure, as the environment becomes hotter. This contrasts with other explanations of unimodality in thermal responses such as enzyme kinetics (***Kontopoulos et al., 2018***; ***Arroyo et al., 2022***) or the metabolic niche hypothesis (***Clarke and Gaston, 2006***) which invoke reduced metabolic rates at high temperatures (above the OTR), either because of the inactivation of enzymes or the reduction in number of viable metabolic strategies, to explain the decline in coexisting species. Furthermore, the observed patterns of negative covariance seen in existing data (analysed here; (***Smith et al., 2019, 2021***)) suggest that peaks of richness should occur towards the higher end of the operational temperature ranges (OTRs) of most mesophilic bacteria, a prediction that is consistent with unimodal microbial species temperature-richness relationships observed in the real world (***Milici et al., 2016***; ***Sharp et al., 2014***; ***Thompson et al., 2017***). We expect that the mechanism we propose here will be particularly relevant to predicting the temperature-richness relationship in: (i) communities where system dynamics is driven primarily by species interactions (as opposed scenarios where environmental filtering or neutral processes dominate); (ii) environments where species typically experience temperatures within their OTR (arguably the most common scenario on planet Earth); (iii) At scales where trait TPC distributions are relatively constant across communities and thus independent of the local environment. At larger scales we expect that processes such as local adaptation are likely to alter these distributions (***Kontopoulos et al., 2018***) as organisms adapt to local temperature regimes. More work is required to test this more explicitly however, and will require datasets explicitly measuring within-community variation of thermal responses across taxa.

We found that the data from the single lab experiment ((***Smith et al., 2021***)) show a greater variance in *E*_*r*_ as well as a stronger covariance between *B*_0,*r*_ and *E*_*r*_ than the literature-synthesised (***Smith et al., 2019***) data (***Figure 4***). This drives a constriction of growth rate variation at ∼ 23°C in the experimental data, which in turn results in a higher predicted peak of species richness at ∼ 20°C these data. Estimates for *E*_*r*_ and *B*_0,*r*_ in both datasets were obtained using comparable methods, so this difference most likely reflects biological and experimental differences between them. Given that the single experimental dataset is for a far more restricted set of thermal taxa from a specific habitat (soils), it is surprising that the TPCs vary more that single community than across the wider diversity of taxa in the literature-synthesised dataset. This either reflects some sort of systematic bias in the literature data, that the local community sampled in the single experiment is a non-random set of co-evolved taxa, or both. In particular, the temperature at which the growth rate variation constricts in the lab dataset is almost identical to the temperature at which those strains were maintained, suggesting a role of species sorting, acclimation or evolution. The literature-synthesised dataset on the other hand represents a much more random set of taxa. Interestingly, the predicted ∼ 9°C peak in species richness based on these data is almost identical to that observed by ***Thompson et al., 2017*** from a wide range of environmental samples, which also presumably emerges from a heterogeneous set of taxa.

We have focused on mathematical feasibility as a constraint on species richness because it is a necessary condition for stable coexistence. We argue that feasibility is a particularly relevant important criterion in microbial communities because stability per se is irrelevant given the continuous assembly and turnover dynamics driven by immigration and emigration that these systems experience. Under such dynamic conditions, richness is arguably constrained heavily by feasibility as only species that are able to enter a system and maintain a non-zero population size will be able to invade. Furthermore, previous work has shown that for feasible fixed points in Lotka-Volterra systems are almost always stable (***Gibbs et al., 2018***).

While our analytical approximation well captures the temperature-richness relationship in statically-assembled synthetic communities (***Figure 2***), it is less accurate in capturing the response in dynamically-assembled ones (***Figure 3***), where the predicted and observed richness values deviate at extreme temperatures. This is likely because during dynamic assembly the distributions of trait values within each community start to deviate, due to species sorting, from those in the invader pool, breaking the assumption we make about the log-normality of the trait values at any given temperature. Additional work that explicitly accounts for the effects of dynamic community assembly on within-community trait distributions will be needed to develop a better approximation (see (***Rossberg, 2013***)). Nevertheless, we emphasise our qualitative predictions remain robust in these dynamically assembled microbial communities. We also expect that while our qualitative predictions will remain robust, the introduction of specific interaction structures will likely alter system dynamics such that the mean-field approximation we use is no longer valid. This is because interaction structure such as modularity (i.e., the tendency of populations to form subsets of stronger interacting “modules”) couples some populations more tightly than others making the overall average interaction a poor approximation of what individual populations experience.

We also note that the Lotka-Volterra model underlying our theory assumes a physically well-mixed system, that is, spatial structure does not play a role. As such, spatial structure will impact species coexistence, for instance, by localising competitive exclusion to spatial “pockets”. We expect that future work incorporating spatial structure in our framework may reveal differences in the thermal responses of microbial species richness between environments with contrasting spatial structures (e.g., soil versus water).

Finally, we acknowledge that we have only considered competitive interactions here. Whilst it has been argued that competitive interactions dominate in microbial communities (***Foster and Bell, 2012***) there has more recently been a recognition of the importance of cooperative interactions that develop through cross-feeding between strains on their metabolic-by-products (***Goldford et al., 2018***; ***Marsland et al., 2019***; ***Lechón et al., 2021***). The generalised Lotka-Volterra model we use is inappropriate for modeling the dynamical consequences of cooperative interactions due to their inherently destabilising effects in such systems due to the absence of explicit resource dynamics (***Bunin, 2017***). Our approach towards determining the temperature dependence of trait distributions could however be applied to other models such as the recently-introduced microbial consumer-resource models (***Marsland et al., 2019***), which would allow explicit characterisation of resource mediated interactions and thus the higher-order interactions and indirect effects that arise. Again, we still expect the broad effects of distributions of thermal response parameters to have similar effects (as the thermal responses of traits is independent of the system dynamics) though the exact mapping of trait distributions (and the traits that need to be considered) on to richness may vary.

Overall, our results provide a compelling theoretical basis and and empirical evidence that the temperature-richness relationship in microbial communities can be strongly driven by variation in thermal physiology across species. Whilst often ignored, quantifying this variation in local communities is likely to be key to predicting the effects of temperature fluctuations on microbial community diversity across space and time.

## Methods

### Derivation of the theory

We begin with the Lotka-Volterra competition model of an *N*-species community where the biomass growth of the *i*th species given by

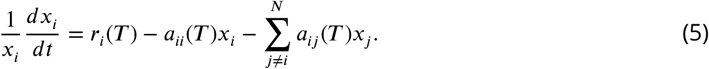

Here, *x*_i_ is its biomass density (abundance) (mass ·volume^−1^), *r*_*i*_(*T*) it’s intrinsic growth rate (time^−1^), *a*_*ij*_ (*T*) is the effect of interaction with the *j*th species’ population (volume·mass^−1^ · time^−1^) (and thus *a*_*ii*_(*T*) is the strength of its intraspecific interactions). All parameters are functions of temperature (*T*), which will omit for now to be specified later.

#### Mean-field approximation of the Lotka-Volterra Model

To determine the feasibility of a community in terms of the parameters in ***Equation 5*** and species richness, we need to first derive an expression for equilibrium biomass, 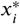. Because it is not possible to meaningfully link the structure of species interactions to the exact closed-form analytical solution for 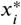 in the Lotka-Volterra model, we use a mean-field approximation (***Wilson et al., 2003***; ***Wilson and Lundberg, 2004***; ***Rossberg, 2013***). By focusing on the averaged effect of interactions on each population’s abundance, this approximation allows us to relate the equilibrium abundance vector to the mean pairwise interaction strengths (*ā*_*ij*_) across the community. We start by rewriting the summed interactions term for the *i*th species in the Lotka-Volterra model as:

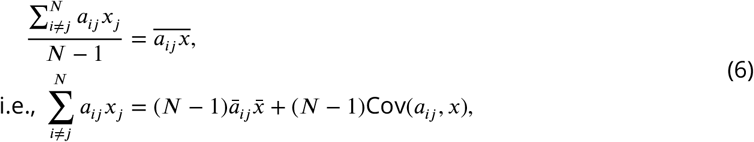

where the bar notation represents the average of the quantity across the *N* − 1 other species that the focal one can interact with (ignoring self-interaction). ***Equation 6*** partitions the effects of interactions on the *i*th species’ population into the average effect, 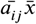, and the covariance between strengths of the interactions and the heterospecifics’ biomasses, cov(*a*_*ij*_, *x*). This mean-field approximation assumes that this second term is negligible, which is equivalent to saying that any individual interaction between the focal species and another species’ population has a small effect on its biomass abundance. We also assume that system (*N*) is large, which ensures that the difference between the average heterospecific’s biomasses and that of the focal species is small (as it is of order *N*^−1^) and can thus be ignored. Combining ***Equation 5*** and ***Equation 6***, we can express each species’ population dynamics in terms the average interaction strength, giving the full mean-field model:

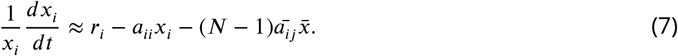

Next, we obtain an expression for the community’s dynamic equilibrium by setting ***Equation 7*** equal to zero and solving for *x*_*i*_, giving:

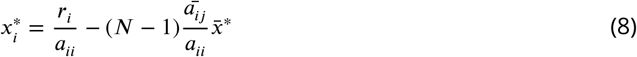

Then, taking the average across the *N* populations and rearranging, the average biomass in the community is:

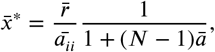

where *ā*_*ii*_ denotes the average intraspecific interaction strength and 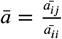 the average of interactions relative to the intraspecific interactions. By expressing interactions relative to the strength of interspecific interactions the new term *ā* measures the effective strength of competition in a community. This aligns with classic results from ecological theory that species coexistence is based on the ratio of inter- and intraspecific competition. We can then substitute the expression for 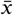 into ***Equation 8*** to get equilibrium biomass:

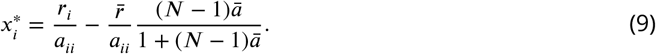

***Equation 9*** shows how the equilbirium abundance reached by a population is a balance between its own growth and intraspecific interaction strength in the first term (which can be shown to be its carrying capacity by setting *a*_*ij*_ = 0 in ***Equation 5***) minus the negative effect of interactions in the second. This second term includes both the average growth-rate across the community as well as a saturating function of interactions. Biologically this makes sense because the effect of competition on a focal species’ biomass depends on the abundance of its competitors in the environment (captured in the 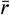 term) and the strength of its interactions with them (captured by 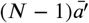). Because interactions are competitive, they will always reduce population biomass relative to intrinsic carrying capacity.

#### Condition for feasibility

Next, we use ***Equation 9*** to derive an expression for community feasibility—which sets the upper bound on species richness *N*—, in terms of species-level traits (i.e., the *r*_*i*_’s and *a*_*ij*_’s). An community is feasible if all its populations have non-zero equilibrium biomasses (i.e.,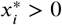), a condition that can be expressed as:

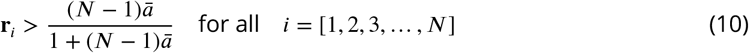

Here, 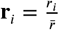 is the relative growth rate of the *i*th species (i.e., its value relative to the average across all *N* populations). ***Equation 10*** states that a community is feasible as long as the the negative effects of competition on each population (RHS) do not outweigh its relative growth rate (LHS).

Using ***Equation 10*** we next derive an expression for *P*_*feas*_, the probability that a *N*-species community is feasible given the distribution of community-level trait values (**r**’s and *a*’s). To do so we treat **r** and *a* in ***Equation 10*** as random variables that follow specific distributions (across species) in the community (denoted by the loss of subscript). This allows us to consider **r**’s cumulative density function (CDF) which gives the probability that any given value of **r** is less than or equal to some value: *F*_**r**_ (*x*) = *P* (**r** ≤ *x*). Because the condition for feasibility states that **r** must be greater than the (negative) effect of interactions, we can use this CDF and the condition in ***Equation 10*** to express *P*_*feas*_ as

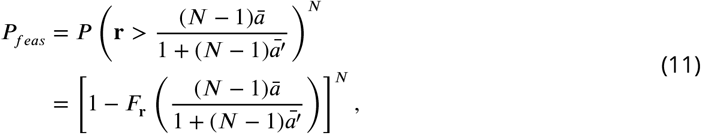

giving the probability of feasibility of an ecosystem as a function of species’ traits. The expression is raised to the *N*^th^ power because all *N* populations within a community must be feasible (the term in the brackets) for a community to be feasible.

#### Incorporating thermal responses of traits

Having defined the relationship between the upper bound of species richness and the distributions of the community-level traits, we now turn to the effect of temperature. For this, we consider how the distribution of **r** and *ā* across populations change with temperature, thus changing *P*_*feas*_ and thus *N*. We derive the distributions of these quantities in terms of the distributions of the thermal response (TPC) parameters of the two underlying population-level traits (growth and interaction rates, i.e., *r*_*i*_’s and *a*_*ij*_’s respectively). For this, we use the Boltzmann-Arrhenius equation to represent the temperature dependence of both these traits (***Gillooly et al., 2001***; ***Savage, 2004***; ***Dell et al., 2011, 2014***):

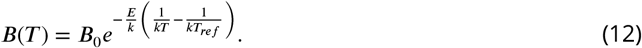

Here, *B*(*T*) is the relevant trait’s value, *T* is temperature in Kelvin, *B*_0_ is the normalisation constant, i.e., the trait value at some reference temperature (*T*_*ref*_, which we set to the middle of the OTR with no loss of generality, we can always obtain the same TPC for a given *T*_*ref*_ by normalising *B*_0_), *E* (eV) is the thermal sensitivity which determines the change in trait value to a unit change of 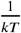,and *k* is the Boltzmann constant. Although species-level thermal performance curves are generally unimodal, the Boltzmann-Arrhenius equation captures the the rising portion (before the temperature of peak performance) of TPCs, which is also the temperature range within which populations typically operate (or experience) (the “Operational Temperature Range”, or OTR; ***Dell et al. (2011***); ***Smith et al. (2019***, 2021)). Indeed, the thermal optima of growth rates of mesophilic prokaryotes in laboratory experiments are typically 5–10 °C higher than their (constant) ambient temperature ***Smith et al. (2019***)). Thus, focusing on the Boltzmann-Arrhenius portion of TPCs is relevant to the dynamics of real microbial communities, and also, conveniently, affords us analytic tractability.

Also, while the validity of ***Equation 12*** for *r* is well established (***Smith et al., 2019***), there are currently no empirical data nor theoretical models that directly support its validity for the temperature dependence of interaction strengths *a*_*ij*_ and *a*_*ii*_ for heterotrophic microbes. Here, we posit that a Boltzmann-Arrhenius (or at least exponential-like) temperature dependence of interaction strength is indeed inevitable because pairwise microbial competitive interactions are ultimately driven by the two species’ resource uptake rates, each of which are known to follow a Boltzmann-Arrhenius form within the OTR (***Smith et al., 2021***; ***Bestion et al., 2018***).

We now consider how the TPC parameters *B*_0_ and *E* of growth (*r*_*i*_’s) and interaction rates (*a*_*ij*_’s) vary across species within the community and how this variation is propagated through ***Equation 12*** to give the community-level distributions of these two traits at different temperatures. We begin with the natural log of ***Equation 12***:

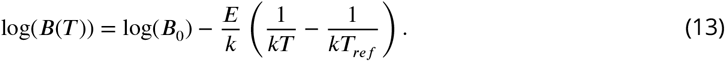

Next, we assume that log(*B*_0_) and *E* are distributed as a multivariate normal distribution such that:

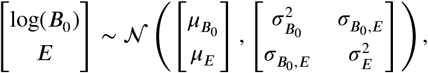

where 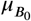 and *μ*_*E*_ are the respective means and 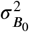 and 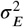 the variances of the normalisation constant and thermal sensitivity respectively, and 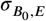 is the covariance between them. *B*_0_ is in-deed expected to be log-normally distributed for growth and interaction rates (***Kontopoulos et al., 2020***; ***Dell et al., 2014***; ***Bestion et al., 2018***). On the other hand, *E* distributions tend to be right-skewed (***Kontopoulos et al., 2020***; ***Smith et al., 2019***; ***Dell et al., 2011***), but we use the normal distribution here as an adequate approximation (see section on numerical evaluation below). Then, because ***Equation 13*** is a linear combination of two co-varying normally-distributed random variables, log(*B*(*T*)) will itself be normally distributed as:

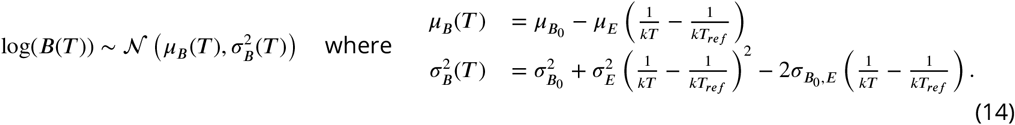

That is, the temperature-specific trait values across species in a community for either growth or interaction rate can be represented by a log-normal distribution, which reveals four key insights (***Figure 1***E). First, the mean trait value across species at a given temperature (*μ*_*B*_(*T*)) increases with their mean baseline trait values 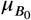s as well as mean thermal sensitivities *μ*_*E*_ s. Note that −*μ*_*E*_ still implies a positive gradient with respect to temperature because we are dealing with inverse temperature (1/*kT*). Second, variation in the trait’s value across species 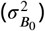 increases with the variance in baseline trait value 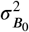. Third, this trait variation decreases to a minimum at some intermediate temperature because the quadratic term 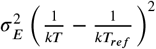 is convex (concave upward) due to the inverse temperature scale. Fourth, the temperature at which this minimum trait variation occurs is modulated by the covariance term 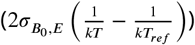. Again noting that we are dealing with inverse temperature here, a negative covariance between the two TPC parameters will increase the temperature of minimum trait variance while a positive covariance will decrease it. This temperature of lowest trait variation is key because it determines the location of the peak of the temperature-richness relationship, as we will show below. Henceforth, we choose *T*_*ref*_ to always be the center of the OTR (∼ 13°C based on our empirical data synthesis; see below). Note that our results are qualitatively independent of our choice of *T*_*ref*_ as one can always recover the same trait-distribution by altering the variance 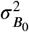 and covariance 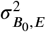 terms.

#### Temperature dependence of species richness

Next, we use (***Equation 14***) to derive the distribution of **r** as well as the value of *ā*, which together determine feasibility (***Equation 11***; ***Figure 1***D). First, recall that:

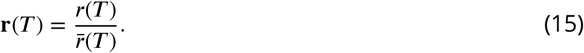

Then, because *r*(*T*)’s TPC follows a Boltzmann-Arrhenius relationship, its TPC parameters are distributed as in ***Equation 14*** and its mean (as a log-normally distributed variable) is given as:

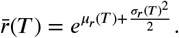

Substituting this into ***Equation 15*** and taking the natural log gives:

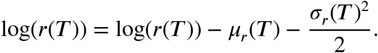

as log(*r*)(*T*) is normally distributed this represents a simple shift in its mean giving ***Equation 3*** of the main text,

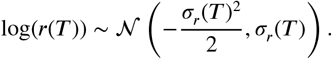

Next consider the thermal dependence of *ā* which depends on the interaction strength (*a*_*ij*_ (*T*)) distribution. Because each *a*_*ij*_ (*T*) also follows a Boltzmann-Arrhenius response, just like **r**, its TPC parameters are also log-normally distributed as in ***Equation 14***. We can therefore obtain its average with the expression (***Equation 4*** in the main text):

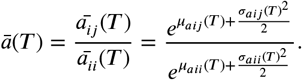

The latter two equations (***Equation 3*** and ***Equation 4*** of the main text) show how the thermal responses of **r** and *ā* are both driven by the variance in the underlying log-trait distribution (and thus the variance in thermal sensitivity 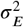 and covariance 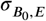) with *ā* additionally being driven by the average log-trait value (and therefore, its average thermal sensitivity, *μ*_*E,a*_). The effects of this on richness are detailed in the main text.

### Numerical Evaluation of the Theory

To illustrate the effect of the community-level distributions of species’ thermal response parameters on richness predicted by our theory, we substitute ***Equation 3 Equation 4*** into the expression for *P*_*feas*_ (***Equation 11***), and numerically solve for *N*. To this end, we set a threshold value of probability of feasibility, *θ* = 0.5, as well as values for the parameters controlling the distributions of the thermal responses for both *a* and **r**. We then calculate the probability of feasibly across a range of *N* values (1 − 1000) using ***Equation 11*** to find the maximum species richness value that has a probability of feasibility greater than the threshold *θ*. Repeating this procedure at different temperatures gives the full temperature-richness curve for any given set of parameters. We calculated such species temperature-richness curves for different combinations of means and variances of thermal sensitivity *E* as well as its covariance 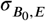 with the normalisation constant *B*_0_ of *r* and *a*.

#### Accuracy of the analytical result

To test the accuracy of our analytical bound on species richness ***Equation 11*** and the effects of temperature on it, we performed two sets of simulations.

First, we obtained species richness values across temperatures by simulating the full GLV model ***Equation 5*** and compared it with the predicted values. For this, we specified the distributions of growth and interaction rates (as defined by ***Equation 14***) to predict richness *N*_*pred*_ as above, with *θ* = 0.5. We then generated 50 replicate GLV communities of sizes *N*_*pred*_ ± 10 parameterised with the same trait distributions, and numerically integrated them to steady state. At each system size we calculated the probability of feasibility (i.e., the proportion of replicates where no extinctions occurred) and found the richness value at which this was the closest to *θ* = 0.5. This gave the system size which in the simulations had the probability of feasibility that matched the given value, and allowed comparison with the predicted values. We carried out this process over a thermal gradient from 0 − 25°*C* and whilst varying the mean, variance and covariance in *E* and *B*_0_ as above.

Second, we simulated the dynamic assembly of replicated GLV communities to allow systems to reach their maximal feasible richness value. For this, we started each such community as an empty system which then was allowed to assemble through sequential species invasions. Each invader has thermal performance traits (*B*_0_s and *E*s) for growth and interactions drawn from a global distribution (i.e., a multivariate log-normal distribution; ***Equation 14*** was sampled to calculate the actual values of the *r*s and *a*s) at that temperature using ***Equation 12***. Following each invasion we numerically integrated the system till it reached steady state, removed extinct populations and record the species richness. We simulated 2000 invasions for each such assembly sequence, which guaranteed that richness reached a quasi-equilibrium where (was relatively stable over time due to a immigration-extinction balance; ***Pawar*** (***2009***)). We varied the three main thermal response parameters (*μ*_*E*_ ∈ [0.0, 0.6], *σ*_*E*_ ∈ [0.01, 0.2] and 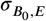 ∈ [0.0, −0.1]) one at a time across the simulations. For each parameter combination, we simulated assembly 20 times over 16 different temperatures ranging from (ranging from 8-22°*C*). To obtain the analytical prediction corresponding to each simulated community, we used the same approach as above: generate predictions of maximal community richness given a distribution of thermal response parameters setting the lower probability threshold as *θ* = 1.0 × 10^−6^. We set *θ* to this value to match the richness predictions at the reference temperature 13°C.

All simulations were carried out using the Julia programming language with the `DifferentialEquations.jl` package for numerical integration (***Rackauckas and Nie, 2017***).

### Empirical data

In order to obtain empirically relevant estimates of the mean, variance and covariance of *B*_0_ and *E* we used data from both ***Smith et al. (2021***) who experimentally measured the thermal performance (growth rate) of 29 strains of environmentally isolated bacteria and ***Smith et al. (2019***) who synthesized data from existing bacterial thermal performance experiments for 422 stains. For both datasets, took the original data and fit the Sharpe Schoolfield model which describes the unimodal thermal response of traits to temperature (including *B*_0_ and *E* values) using the rTPC package (***Schoolfield et al., 1981***; ***Padfield et al., 2021***). We rejected any fits that had non-significant (*p* < 0.05) parameter estimates or did not converge. Taking the fitted *B*_0_ and *E* values, normalised the *B*_0_ values by dividing by the mean to allow comparison across the datasets, and filtered our the values of log(*B*_0_) larger than −15. We then fitted the multivariate-normal distribution using maximum likelihood estimation (MLE; ***Besançon et al. (2021***)) giving estimates for the means and variance-covariance matrix, which can be used to generate temperature dependent distributions of growth rate across the community ***Equation 14***. We used these parameters to estimate temperature-richness relationships using the method described in the previous section with both *r* and *a* TPC parameters set to the same values except for the 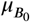 values which were set to 0.0 and −5.0 for log(*r*_0_) and log(*a*_0_) respectively.

